# Clonal cell states link Barrett’s esophagus and esophageal adenocarcinoma

**DOI:** 10.1101/2023.01.26.525564

**Authors:** Rodrigo A. Gier, Raúl A. Reyes Hueros, Jiazhen Rong, Maureen DeMarshall, Tatiana A. Karakasheva, Amanda B. Muir, Gary W. Falk, Nancy R. Zhang, Sydney M. Shaffer

## Abstract

Barrett’s esophagus is a common type of metaplasia and a precursor of esophageal adenocarcinoma. However, the cell states and lineage connections underlying the origin, maintenance, and progression of Barrett’s esophagus have not been resolved in humans. To address this, we performed single-cell lineage tracing and transcriptional profiling of patient cells isolated from metaplastic and healthy tissue. Our analysis revealed discrete lineages in Barrett’s esophagus, normal esophagus, and gastric cardia. Transitional basal progenitor cells of the gastroesophageal junction were unexpectedly related to both esophagus and gastric cardia cells. Barrett’s esophagus was polyclonal, with lineages that contained all progenitor and differentiated cell types. In contrast, precancerous dysplastic foci were initiated by the expansion of a single molecularly aberrant Barrett’s esophagus clone. Together, these findings provide a comprehensive view of the cell dynamics of Barrett’s esophagus, linking cell states along the full disease trajectory, from its origin to cancer.

## Introduction

Metaplasia is a response to injury in which the cell types normally found in a tissue are replaced by other, foreign cell types^1^. Because the cells of a metaplastic tissue take on a new identity, it is difficult to determine their previous identity and tissue of origin. Barrett’s esophagus is a classic example of metaplasia: chronic exposure to acid reflux is believed to cause the squamous cells of the esophagus to be replaced by columnar cells resembling those of the stomach and the intestine^2^. The cell of origin for Barrett’s esophagus and whether it arises through the expansion of a single cell or multiple cells remains highly debated^3^. It is also unclear whether a single stem cell can generate all of the cell types found in Barrett’s esophagus, or if different cells contribute to separate compartments within the tissue^4^. As with other metaplastic tissues, Barrett’s esophagus can undergo additional changes that lead to cancer^5^; however, the early molecular events that initiate this malignant transformation and the specific cell types involved are not fully understood^6^. Hence, to capture the origin of Barrett’s esophagus and its progression to cancer, we need detailed tracking of cell lineages within these tissues to build a trajectory of how cells advance through the stages of disease.

As applied to the origin of Barrett’s esophagus, reporter-based mouse models for lineage tracing have shown that metaplasia can develop from several different types of cells, including progenitors in the gastric cardia^7^, the esophagus^8^, and the gastroesophageal junction itself^9,10^. However, it is not possible to use this lineage tracing approach in humans, since it requires genetic engineering to express lineage tags. Thus, lineage tracing in human tissue has largely relied on sequencing for the detection of somatic mutations present in bulk samples, which lacks the resolution to attribute mutations to individual cells^11^. Alternatively, individual crypts or single cells can be isolated with laser capture microdissection and then sequenced for somatic mutations, but this approach is not feasible for large cell numbers^12,13^. In both of these techniques, identical mutations detected in separate samples suggest a common origin for the cells within them. However, the identity of these cells remains unresolved, making it impossible to precisely determine the cell types that share a common origin with Barrett’s esophagus.

Lineage tracing of esophageal adenocarcinoma development has similarly relied on bulk sequencing of mutations in the cancer and adjacent Barrett’s esophagus tissue^14–17^. In line with our understanding of cancer evolution^18^, esophageal adenocarcinoma is initiated by a clonal expansion, and a recent transcriptional analysis suggests that it originates from metaplastic cells^11^. However, the cancer often shares minimal mutational overlap with the adjacent Barrett’s esophagus, suggesting that bulk lineage tracing is failing to capture the cells that initially transform, since they represent a small subset of these bulk samples. Furthermore, despite extensive genetic characterization of esophageal adenocarcinoma, the drivers of this transformation to cancer are still largely unknown. Hence, to identify these early molecular changes, we need single-cell resolution to track the malignant transition of a Barrett’s esophagus cell.

To address the gaps in our knowledge of how Barrett’s esophagus develops from normal cells and progresses to esophageal adenocarcinoma, the challenges are two-fold: we must have single-cell lineage tracing to identify how cells are related to each other, and we must also know the transcriptional states within these cells. To overcome these challenges, recent advances in single-cell sequencing have now enabled the detection of single-cell lineage information–in the form of mitochondrial DNA (mtDNA) mutations–with high-throughput single-cell RNA sequencing (scRNA-seq)^19,20^. Because mtDNA is passed on through cell division, each mutation labels cells that are derived from a common progenitor cell. By linking cells with the same mtDNA mutations, we can reconstruct their lineages. Beyond determining lineage relationships between individual cells, pairing lineage and cell state information allows us to directly measure how changes in gene expression determine cell fate.

In this study, we seek to elucidate the full trajectory of Barrett’s esophagus disease by applying single-cell lineage tracing paired with transcriptomics to patient samples from the gastroesophageal junction. We also apply this lineage-based approach to precancerous dysplastic samples, allowing us to directly resolve the molecular factors and cell dynamics that promote cancer. We show that mtDNA mutations resolve clones that span tissue types as well as different stages of disease. We also find that lineages within Barrett’s esophagus are maintained by stem cells that give rise to all differentiated cell types, including newly discovered rare populations. Finally, clonally heterogeneous Barrett’s esophagus evolves into esophageal adenocarcinoma through the expansion of a single clone whose aberrant cell states generate a competitive advantage.

## Results

### Transitional basal progenitor cells are related to the squamous esophagus and gastric cardia

The tissues spanning the gastroesophageal junction (Fig. 1A) are maintained by progenitor populations that have the potential to become dysregulated in response to chronic acid reflux^21^, and thus, could be the cell of origin for Barrett’s esophagus. Moreover, recent profiling of Barrett’s esophagus by scRNA-seq found important transcriptional similarities between Barrett’s esophagus and neighboring normal cell types from esophageal submucosal glands and the gastric cardia^11,22^. However, from transcriptional states alone it is impossible to know whether single cells are indeed related.

**Fig. 1.**
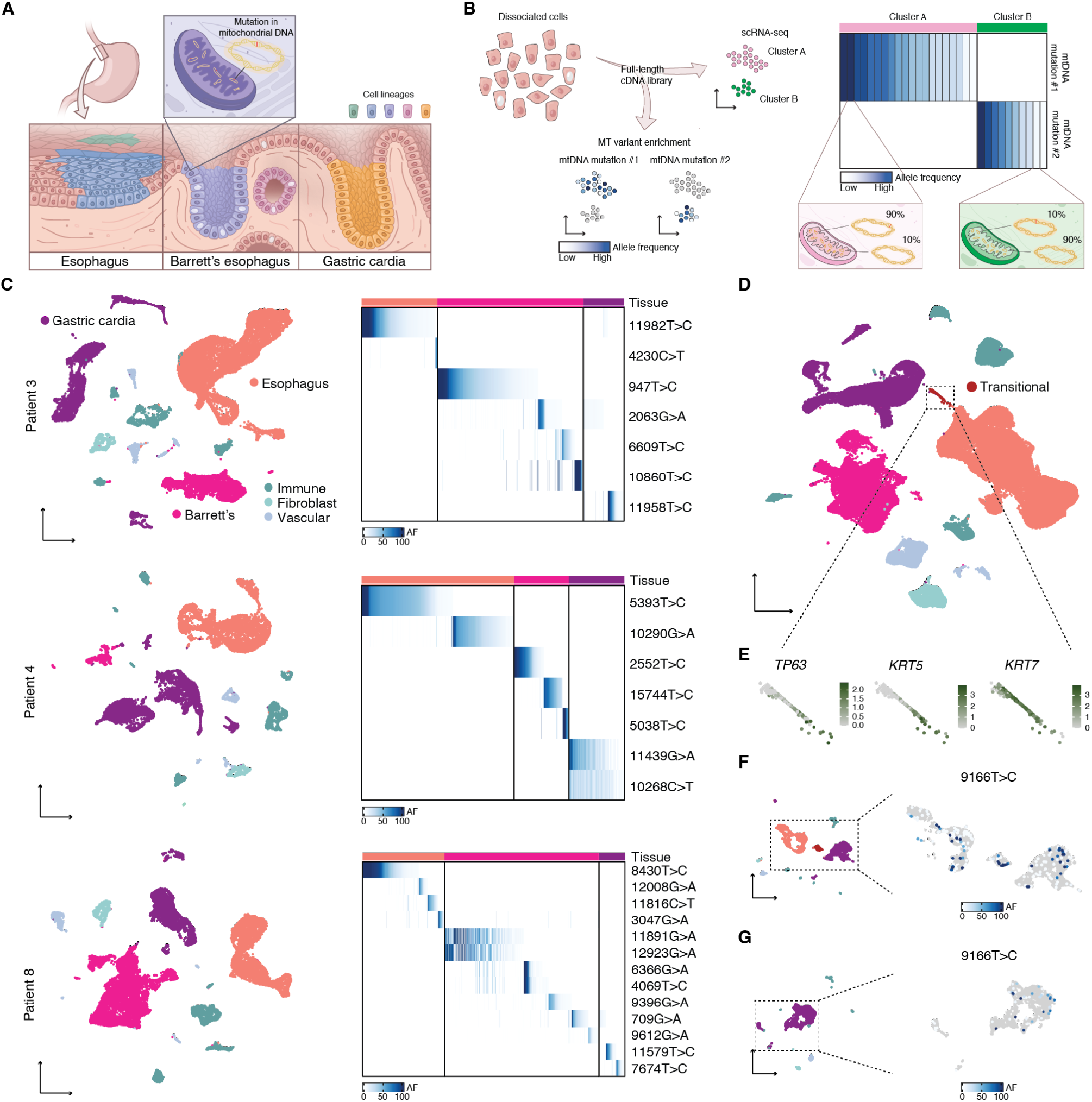
Cell lineages in human gastroesophageal junction tissues labeled by mtDNA mutations. (**A**) Barrett’s esophagus occurs at the gastroesophageal junction between esophageal squamous and gastric cardia tissues. Clones within these tissues can be traced using distinctive mtDNA mutations. (**B**) Conventional scRNA-seq libraries can be enriched for mtDNA mutations, enabling the linking of clones to cell states. (**C**) UMAPs of scRNA-seq of matching Barrett’s esophagus, esophageal squamous, and gastric cardia tissues for three Barrett’s esophagus patients; tissues contained supporting immune, fibroblast, and vascular cells. Adjacent heatmaps show the allele frequencies (AF) of mtDNA mutations within Barrett’s esophagus, esophageal, and gastric cardia cells. (**D**) UMAP of scRNA-seq of all the samples collected in this study; highlighted are transitional basal progenitor cells from the normal squamocolumnar junction. (**E**) Callouts of the transitional cells from (D) featuring the expression of basal progenitor markers. (**F**) UMAP of scRNA-seq of a single squamocolumnar junction biopsy from patient 9; in the callout is the same UMAP, colored with the allele frequency of mutation 9166T>C. (**G**) UMAP of scRNA-seq of a single gastric cardia biopsy from patient 9; in the callout is the same UMAP colored with the allele frequency of mutation 9166T>C.

In order to determine whether any of these cell types native to the gastroesophageal junction were the source of Barrett’s esophagus in humans, we collected a full set of pinch biopsies of the esophagus, gastric cardia, and Barrett’s esophagus from three Barrett’s esophagus patients at endoscopy and immediately subjected them to scRNA-seq and mitochondrial variant enrichment (Fig. 1B). The different tissue types were readily identifiable after uniform manifold approximation and projection (UMAP) and Louvain clustering, with confirmation of expected cell types by established marker genes (Fig. 1C and Fig. S1). Mitochondrial variant enrichment revealed somatic lineages unique to each tissue, but none that were present across multiple tissues in any of the patients (Fig. 1C). As expected, we found that the esophagus and gastric cardia were polyclonal, which is a well-known feature of normal healthy tissue^23^. However, the absence of a founder clone in any of the Barrett’s esophagus biopsies suggests that Barrett’s esophagus also originates from multiple clones, challenging the current model that posits an initial clonal sweep^16,24^.

Within each biopsy, we found a separate set of discrete lineages, marked by unique mtDNA mutations. We therefore reasoned that each lineage occupied a small space within the tissue and that cells from the same lineage did not migrate far from one another. Thus, it is unlikely that biopsies taken far apart (such as our normal biopsies and the Barrett’s esophagus biopsies) would share a common mutation. To look for relationships between tissues that are spatially close to each other, we next profiled a biopsy obtained from the gastroesophageal junction of a patient with Barrett’s esophagus. Across our entire scRNA-seq dataset of 101,000 cells collected from nine patients, we observed large clusters corresponding to normal esophagus and gastric cardia. However, the sample from the gastroesophageal junction contained a small subset of cells bridging the esophageal squamous and gastric cardia populations (Fig. 1D). We found that these cells expressed *TP63*, *KRT5*, and *KRT7*, which are all markers of a recently discovered transitional basal progenitor cell that was shown to exist natively at the squamocolumnar junction and generate Barrett’s esophagus-like tissue^10^ (Fig. 1E).

Despite their transcriptional similarity, the lineage relationship between transitional basal progenitor cells and neighboring esophageal squamous and gastric cardia cells was not known in humans. Mitochondrial variant enrichment unexpectedly revealed that a shared mutation, 9166T>C, linked the transitional basal progenitor cells to both the esophageal squamous and gastric cardia cells captured within the same biopsy (Fig. 1F). We also detected the same mutation in a separate gastric cardia biopsy from the same patient (Fig. 1G), but not in the biopsy taken from the esophagus, or in any of the other 13 biopsies within our mtDNA mutation dataset. Thus, we can conclude that squamous esophagus, gastric cardia, and transitional basal progenitor cells were directly related, sharing a common progenitor.

### Barrett’s esophagus is a polyclonal tissue whose multiplicity of cell types arises from a single stem cell

In order to understand the role of Barrett’s esophagus in the development of cancer, we first needed to characterize its cell types and clonal structure. In particular, we wanted to know whether a single stem cell could generate all the cell types found in Barrett’s esophagus, or if different progenitor populations contributed to separate compartments within the tissue. We collected Barrett’s esophagus biopsies from seven patients for scRNA-seq and mitochondrial variant enrichment. Of these patients, three were previously diagnosed with various degrees of dysplasia, but dysplasia was not detected in the remaining four patients. When we merged the individual Barrett’s esophagus transcriptomes from all the patients, including those with dysplasia, a set of consensus Barrett’s esophagus cell types emerged from the overlap between known non-dysplastic samples (Fig. 2A and Fig. 2B).

**Fig. 2.**
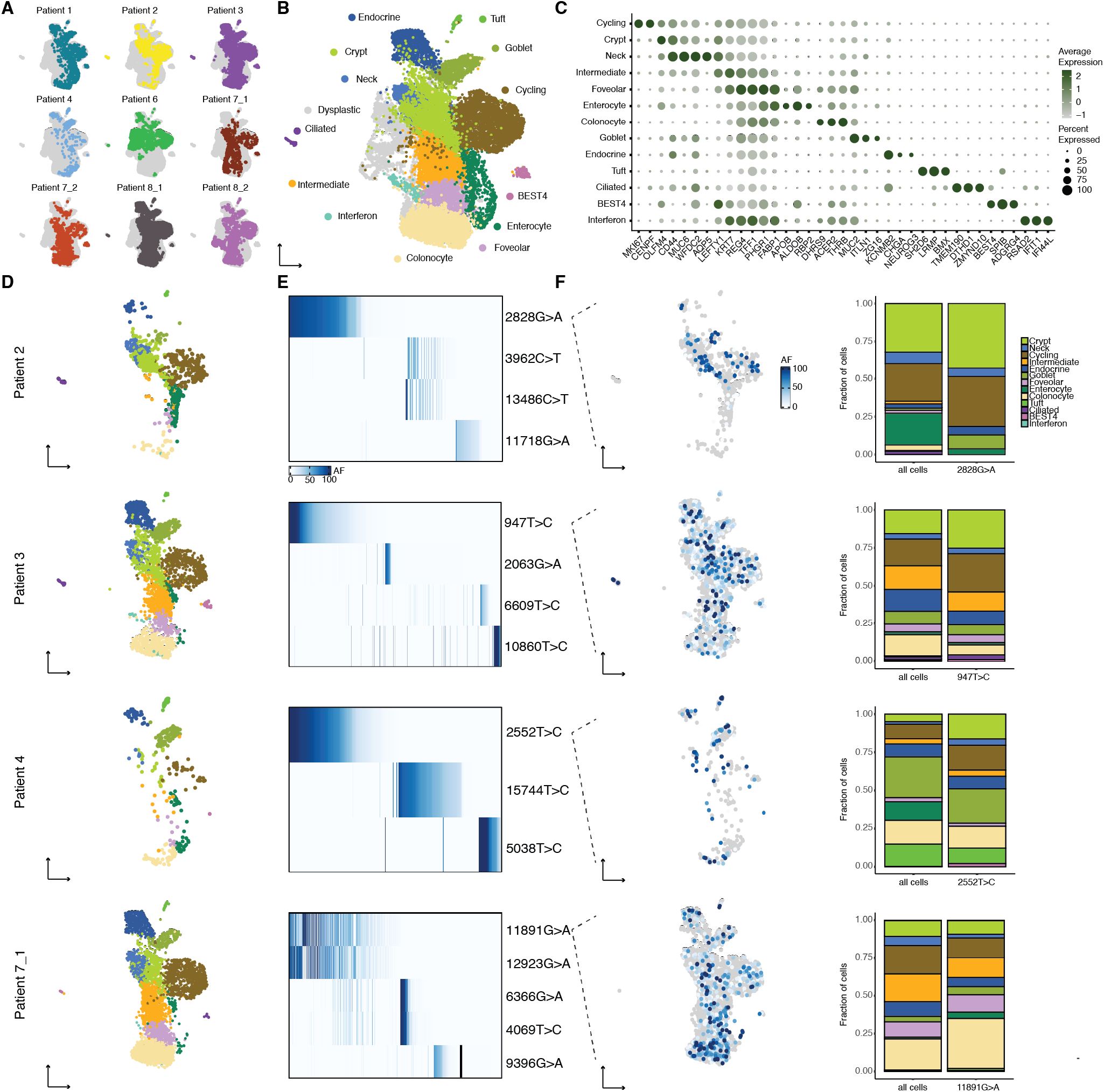
Identity and clonal relationships of Barrett’s esophagus cell types. (**A**) UMAP of scRNA-seq of all Barrett’s esophagus biopsies, with the cells from each individual biopsy highlighted separately. (**B**) UMAP of scRNA-seq of all Barrett’s esophagus biopsies, colored by consensus Barrett’s esophagus cell types common to a majority of non-dysplastic Barrett’s esophagus samples. (**C**) Bubble plot of marker genes for Barrett’s esophagus cell types. (**D**) UMAPs generated from (B) containing only the cells corresponding to the non-dysplastic Barrett’s esophagus sample indicated. (**E**) Heatmaps show the allele frequencies of mtDNA mutations within the Barrett’s esophagus cells in the adjacent UMAP. (**F**) UMAP from (D) colored with the allele frequency of a representative mtDNA mutation; stacked bar graph comparing the proportion of Barrett’s esophagus cell types within the representative lineage to their proportion within the entire sample.

Established Barrett’s esophagus cell types such as mature secretory (goblet and endocrine) and absorptive (enterocyte) cells, as well as OLFM4 stem cells, were well represented in our data. We also discovered new cell types that included tuft cells, airway-like ciliated cells, and poorly differentiated BEST4 cells, all clearly identifiable by distinct markers known from other tissues^25,26^ (Fig. 2B, Fig. 2C, and Fig. S2). Furthermore, the majority of Barrett’s esophagus cell types were present in all four non-dysplastic Barrett’s esophagus biopsies, and we detected all of the cell types in at least three of the biopsies, confirming their fundamental relevance to the tissue (Fig. 2D).

As noted earlier, we did not observe a founder clone within any of the Barrett’s esophagus biopsies, which instead consisted of multiple discrete lineages that accounted for smaller subpopulations of cells (Fig. 2E). Upon overlaying the lineages on our annotated UMAPs, we discovered that each lineage largely resembled the sample as a whole, both in the cell types present, as well as their relative proportion (Fig. 2F). Because each lineage contained all or most of the cell types within Barrett’s esophagus, we reasoned that there must be a single Barrett’s esophagus stem cell that could give rise to the entire multiplicity of Barrett’s esophagus cell types. Such remarkable differentiation potential was supported by the existence of a large pool of immature progenitor cells (Fig. S2).

While histological analysis and protein staining have been used to identify some of the cell types in Barrett’s esophagus, there is currently no comprehensive mapping of the localization of these cell types based upon gene expression within the tissue. We performed spatial transcriptomics using multiplexed hybridization chain reaction^27^ to probe a panel of cell-type specific markers within the same tissue section. We found regular expression patterns along the structure of the glands, with *ALDOB* and *MUC5AC* at the lumen, *MUC2*, *CHGA*, and *NEUROG3* in the middle of the gland, and stem cell markers *LGR5* and *OLFM4* at the base of glands (Fig. 3A, Fig. 3B, and Fig. S3). We also directly confirmed the presence of newly identified cell types, including tuft (*SH2D6*), ciliated (*ZMYND10*), and BEST4 cells (Fig. 3C and Fig. 3D), and found that the ciliated cells specifically localized to the boundary of the squamous esophagus and Barrett’s glands, suggesting a role in the junction of these tissues.

**Fig. 3.**
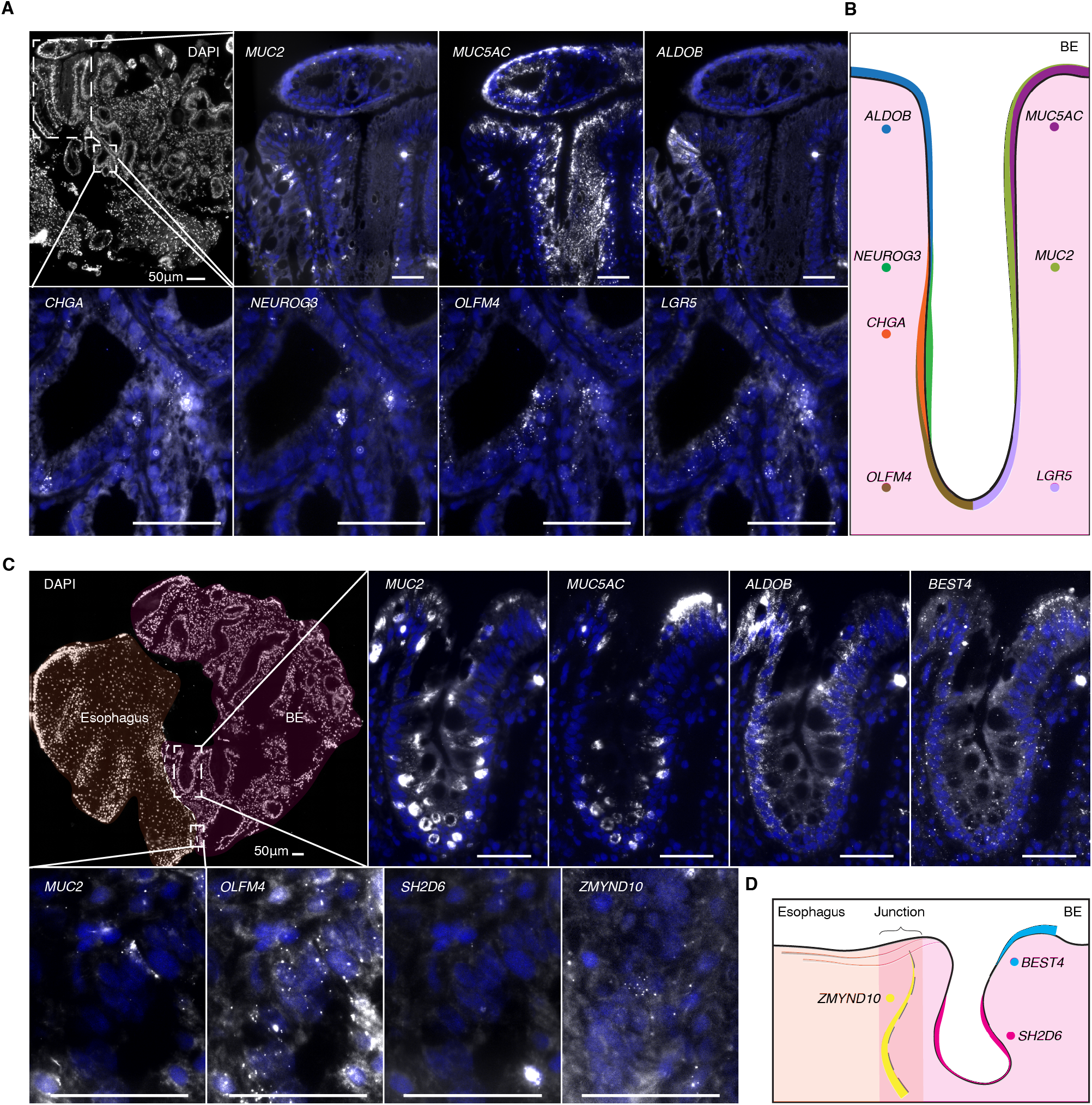
Spatial location of Barrett’s esophagus cell types. (**A**) Multiplexed HCR RNA FISH of fresh frozen Barrett’s esophagus sections for *MUC2* (goblet), *MUC5AC* (foveolar), *ALDOB* (enterocyte), *CHGA* (enteroendocrine), *NEUROG3* (enteroendocrine progenitor), *OLFM4* (stem), and *LGR5* (stem) with DAPI counterstain. Scale bars, 50 μm. DAPI staining is displayed in blue. All genes were detected in sections from at least two patients. (**B**) Schematic depicting the general location of key marker genes within Barrett’s esophagus glands. (**C**) Multiplexed HCR RNA FISH of fresh frozen tissue sections from the junction of Barrett’s esophagus and squamous esophagus for markers of newly discovered rare cell types, as well as typical marker genes from (A): *SH2D6* (tuft), *ZMYND10* (ciliated), and *BEST4*. Scale bars, 50 μm. DAPI staining is displayed in blue. All genes were detected in sections from at least two patients, with the exception of *ZMYND10*. (**D**) Schematic depicting the general location of rare-cell markers within Barrett’s esophagus tissue.

### A single molecularly aberrant stem cell can initiate the malignant transformation of Barrett’s esophagus to esophageal adenocarcinoma

Given that Barrett’s esophagus contained multiple lineages, we next wondered whether one lineage or many lineages were represented in the transition to cancer. Within a biopsy collected from a patient with high-grade dysplasia, we observed two large clusters of epithelial cells by scRNA-seq (Fig. 4A); one of the clusters consisted of cells that expressed established Barrett’s esophagus cell type markers, whereas the other did not, instead showing a marked loss of *PIGR* and *FAM3D*, known regulators of intestinal barrier integrity^28,29^ (Fig. S4). Thus, we concluded that we had captured adjacent patches of non-dysplastic Barrett’s esophagus and dysplastic tissue. In order to understand whether the cells in these molecularly distinct tissues were clonally related, we subjected them to mitochondrial variant enrichment. Consistent with our earlier lineage analysis of Barrett’s esophagus, the Barrett’s esophagus tissue in this sample was made up of several distinct clones, each of which accounted for a fraction of the population (Fig. 4B). The dysplastic tissue, on the other hand, very strikingly originated from the expansion of a single cell containing mtDNA mutation 15153G>A (Fig. 4C). We developed a zero-inflated beta binomial (ZIBB) model that identified a subset of non-dysplastic Barrett’s esophagus cells belonging to the same lineage as the dysplastic cells, confirming that dysplasia originated from Barrett’s esophagus (Fig. S5). Within this dominant lineage, we further observed the emergence of a large dysplastic subclone that was spatially restricted within UMAP space (Fig. 4C). Such clonal expansion was not unique to the one dysplastic case; a second patient with low-grade dysplasia contained a similar precancerous outgrowth (Fig. S6).

**Fig. 4.**
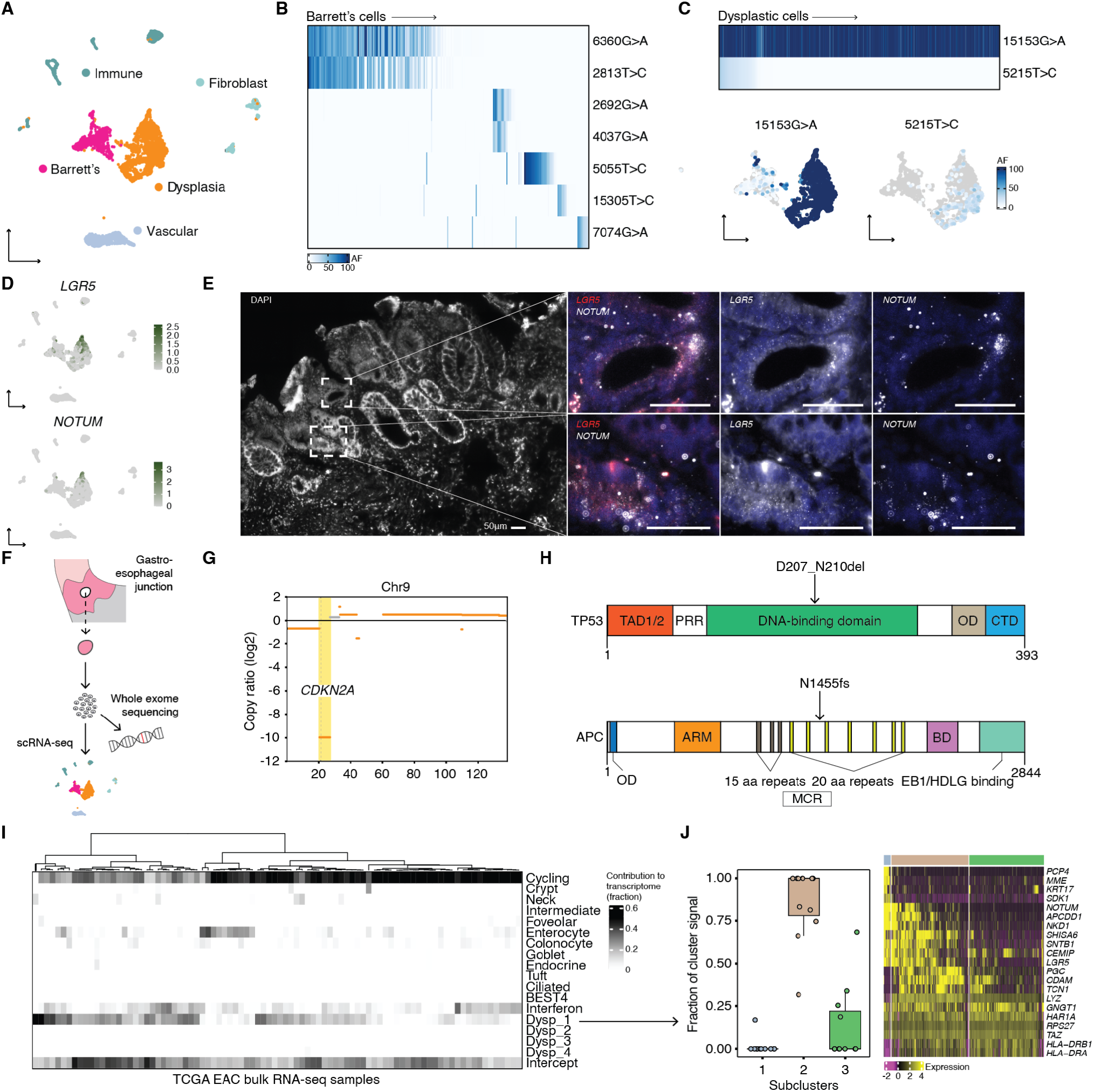
Clonal and molecular characterization of dysplasia and its similarity to esophageal adenocarcinoma. (**A**) UMAP of scRNA-seq of a single Barrett’s esophagus biopsy from patient 6 containing Barrett’s esophagus and dysplastic cells. (**B**) Heatmap shows the allele frequencies of mtDNA mutations within the Barrett’s esophagus cells in the adjacent UMAP. (**C**) Heatmap shows the allele frequencies of mtDNA mutations within the dysplastic cells in the adjacent UMAP; UMAPs of the Barrett’s esophagus and dysplastic cells in (A) colored by the allele frequencies of the dysplastic mtDNA mutations. (**D**) UMAPs from (A) featuring the expression of *LGR5* and *NOTUM*. (**E**) RNA FISH HCR probing for *LGR5* and *NOTUM* in fresh frozen tissue sections of a Barrett’s esophagus biopsy from patient 6. (**F**) Schematic illustrating the sequencing workflow for the cells isolated from a single Barrett’s esophagus biopsy taken from patient 6. (**G**) Copy number analysis from whole exome sequencing of the same cell population from patient 6 that underwent scRNA-seq; featured is chromosome 9, highlighting the loss of *CKDN2A*. (**H**) Schematics of mutations detected in TP53^54^ and APC^55^ proteins by whole exome sequencing. (**I**) Analysis of the contribution of mRNA signals from Barrett’s esophagus and dysplastic cell states to the bulk transcriptomes of esophageal adenocarcinoma tumors from TCGA. Dysp_1 is the set of mRNA signals from the dysplastic cells in (A); Dysp_2-Dysp_4 correspond to mRNA signals specific to the remaining three dysplastic biopsies. (**J**) There were three cell states (or subclusters) within the dysplastic tissue from patient 6. For the cases where at least 3/10 of the bulk transcriptome was contributed by mRNA signals from the dysplastic tissue from patient 6, highlighted is the fraction of that dysplastic signal that each of three cell states accounted for. The heatmap shows genes that were differentially expressed within the three cell states.

Having resolved the clonal structure of the dysplastic tissue and determined its relationship with adjacent non-dysplastic cells, we next investigated whether its transcriptional states could explain how it evolved. We noticed a large increase in the abundance of LGR5 stem cells, which is a known feature of dysplastic progression^30,31^ (Fig. 4D). More surprising was the expression of *NOTUM*, as well as other WNT antagonists^32–35^, within a subset of the dysplastic cells (Fig. 4D). In situ hybridization of tissue from the same patient confirmed the scRNA-seq data and showed an expansion of the stem cell compartment (Fig. 4E). Interestingly, several genes, including *COL17A1*, which drives stem cell fitness in the skin^36^, were differentially expressed in the dysplastic subclone, suggesting that new cell states that emerge within an already transformed clonal population could confer additional malignant potential (Fig. S7).

Given the level of WNT pathway dysregulation, we wondered whether there might be an underlying genetic mutation in a WNT regulator. Thus, we performed whole exome sequencing on cells that were dissociated from the same biopsy that underwent scRNA-seq, along with matching normal squamous esophagus and gastric cardia controls (Fig. 4F). The whole exome sequencing revealed mutations in *CDKN2A* and *TP53*, the genes most frequently implicated in dysplasia and esophageal adenocarcinoma^37,38^ (Fig. 4G and Fig. 4H). Additionally, as we hypothesized, there was a truncating mutation in the mutation cluster region of *APC*, a negative WNT regulator (Fig. 4H). This result suggests that the dysplastic cells were being driven by a mechanism recently discovered in premalignant colon adenomas, where APC-mutant stem cells secrete NOTUM, and thereby shut down and outcompete neighboring wild-type ones^39^.

Next, we wanted to determine whether esophageal adenocarcinoma contained Barrett’s esophagus and dysplastic cell states from our scRNA-seq analysis that would further demonstrate their involvement in its development. We checked for the presence of mRNA signals^40^ corresponding to the Barrett’s esophagus and dysplastic cell states in 88 bulk esophageal adenocarcinoma tumor transcriptomes previously analyzed by TCGA (The Cancer Genome Atlas)^38^ (Fig. 4I). In 11 of the 88 esophageal adenocarcinoma tumors, we found that cell states from the dysplastic sample contributed to at least 3/10 of the bulk transcriptome (Fig. 4I). Specifically, of the three cell states that made up that dysplastic sample, the second accounted for the majority of the signal in the cases where there was significant mapping to a bulk transcriptome, and it included cells with high *LGR5* and *NOTUM* expression (Fig. 4J). Therefore, in this subset of esophageal adenocarcinomas, we can conclude that there is a WNT-dysregulated transcriptional state that is similar to the cell state that we discovered in dysplasia. In other words, Barrett’s esophagus can transition to esophageal adenocarcinoma directly through the dysplastic cell states that we identified.

## Discussion

In this study, we applied lineage-resolved scRNA-seq to human samples collected from the gastroesophageal junction of patients with Barrett’s esophagus and dysplasia. We discovered new biology at every stage of the disease: transitional basal progenitor cells found at the squamocolumnar junction were directly related to both squamous esophagus and gastric cardia cells; Barrett’s esophagus is a polyclonal tissue in which a single stem cell can generate a previously unknown number of mature, specialized cell types; and Barrett’s esophagus can transition to esophageal adenocarcinoma through the expansion of a WNT-activated clone that outcompetes normal neighbors with an inhibitory mechanism described in the development of colon cancer^39^.

While scRNA-seq has proven to be an indispensable tool for mapping normal and diseased human cell states, it also has important limitations. How these cell states are related, for example, cannot be conclusively answered without accompanying lineage information. Our results highlight the value of such information to connect cell states not just within a single phenotypic state, such as one tissue type or disease, but also between phenotypes, connecting different stages of a disease or multiple tissue types through lineage. Without lineage tracing, the scRNA-seq data alone could not have captured the relationship between transitional basal progenitors and esophageal squamous and gastric cardia tissues, as was missed in previous scRNA-seq work that specifically profiled the normal squamocolumnar junction^11^. Similarly, combining lineage tracing with scRNA-seq for different stages of disease allowed us to show that the transformation of Barrett’s esophagus to cancer was caused by a cascade of clonal expansions with uniquely aberrant cell states.

While our study achieves single-cell resolution for the lineage tracing, the resolution of the lineages themselves remains a limitation. Because we are using mtDNA mutations as a marker of cell lineage, we depend upon the acquisition of mutations to provide the lineage resolution. It is thus important to consider that acquisition of mtDNA mutations is not uniform across cells, and is dependent upon the number of mtDNA per cell and the mutation rate, both of which can vary across cell types^41^ and change in the setting of cancer^42^. It is also possible that the mtDNA mutations used for lineage tracing could have pathogenic effects that change their distribution in the tissue. In the future, we expect to see new single-cell methods for lineage tracing in tissues that will improve upon these resolution constraints.

The origins of Barrett’s esophagus and its transformation to esophageal adenocarcinoma are complex phenomena. We outline a mechanism whereby Barrett’s esophagus transforms through a single clone with a WNT-dysregulated cell state. In our TCGA analysis, we confirm that this cell state is relevant to esophageal adenocarcinoma, but in a subset of the cases. While this result represents a pathway for progression in metaplasia, it is likely one of many. Thus, this work further supports a growing appreciation for the fact that the paths between related phenotypic states (such as Barrett’s esophagus to cancer) will not always follow a single trajectory^43^. Studies like ours, even when they profile larger numbers of human samples, will never be exhaustive. Their value lies in providing hard evidence for the role of specific cell states in maintaining and connecting phenotypes that before we could only speculate about. In the case of Barrett’s esophagus, these relationships have important clinical implications that earlier methods struggled to address. We hope that this work further demonstrates the general usefulness of tracking clones across diverse cell states in disease, and justifies its broader application not just at the gastroesophageal junction, but throughout the human body.

## Methods

No statistical methods were used to predetermine sample size. The experiments were not randomized and the investigators were not blinded to allocation during experiments and outcome assessment.

### Patient samples

All samples were obtained following written informed consent under IRB approval (Protocol #813841). Patients had a prior history of Barrett’s esophagus and were undergoing routine surveillance endoscopy. Pinch biopsies were taken from the BE tissue, as well as the normal esophagus proximal to the Barrett’s esophagus segment and the gastric cardia at the discretion of the endoscopist (G.W.F). Samples were immediately transferred for processing in ice-cold Dulbecco’s phosphate buffered saline (DPBS; Corning, 21-031-CV). The Barrett’s esophagus biopsy from patient 2 was the only exception to the above; the tissue was immediately preserved in 1 mL CryoStor CS10 (Biolife Solutions, 210373) and frozen at −80 °C in an isopropyl alcohol-filled freezing container before being transferred to liquid nitrogen after 24 hours.

### Isolation of single cells from patient tissue

Fresh pinch biopsies were washed twice in DPBS before being coarsely chopped with a scalpel and transferred into 0.5 mL DPBS solution containing 0.1 WU ml^−1^ Liberase TH (Roche, 5401151001) and 0.5 U ml^−1^ RQ1 DNase I (Promega, M6101). The samples were incubated at 37 °C in successive 10-minute rounds with gentle vortexing every 2 minutes and trituration every 5 minutes. At the end of each round of incubation, large tissue pieces were spun down in a microcentrifuge and the supernatant containing single cells was passed through a 70-um cell strainer fitted onto a 50-mL conical tube containing 8 mL DPBS + 1% bovine serum albumin (BSA; Sigma-Aldrich, A7906) in DPBS on ice. 0.5 mL fresh dissociation solution was added to the tissue at the beginning of each incubation step. The samples were further filtered through the 0.35 um cell-strainer cap of a FACS tube and single cells were pelleted by centrifugation at 400g for 4 minutes at 4 °C. Samples were washed twice in 1 mL DPBS + 0.04% BSA and finally resuspended in 50 uL DPBS + 0.04% BSA, at which point they were counted in 0.2% trypan blue (Gibco, 15250061) in preparation for 10x Genomics Chromium Next GEM Single Cell 3’ v3.1 library preparation. For the frozen BE biopsy from patient 2, the tissue was quickly thawed at 37 °C and rinsed three times in DPBS + 10% fetal bovine serum (GE, SH30396.03) before being dissociated and filtered as described above. After the final filtering step, dead cells were removed with the Dead Cell Removal Kit (Miltenyi Biotec, 130-090-101) following the manufacturer’s protocol.

### 10x Genomics Chromium Next GEM Single Cell 3’ v3.1 library preparation and sequencing

Single-cell suspensions were loaded onto a 10x Chromium Controller for GEM generation of approximately 5,000-10,000 cells for each sample. 10x Genomics Chromium Next GEM Single Cell 3’ v3.1 Dual Index library preparation was performed following the manufacturer’s protocol. Library quality was confirmed on an Agilent 2100 Bioanalyzer Instrument using the High Sensitivity DNA Kit (Agilent). Libraries were paired-end sequenced on an Illumina NextSeq 550 with 28 cycles for Read 1 and 43 cycles for Read 2, as well as 10 cycles for both indices.

### 10x Genomics sequencing data mapping and count matrix generation

Raw Illumina base call files were demultiplexed and converted into FASTQ files using CellRanger mkfastq v5.0.0. The resulting FASTQ files were loaded into STARsolo v2.7.9a for alignment to the 10x reference GRCh38-2020-A^44^. Count matrices generated with the -soloFeatures GeneFull argument were used for downstream analyses.

### scRNA-seq dimensionality reduction, clustering, and cell-type annotation

Count matrices for each sample were converted into Seurat objects in R (Seurat, v4.0.2) and genes present in fewer than three cells were removed^45^. Doublets were detected using scDblFinder (v1.8.0), with nfeatures = 3000 and includePCs = 1:20^46^. Once doublets had been filtered out, only cells with more than 300 genes and less than 30% mitochondrial reads were retained. Outlier cells containing high total counts were also removed on a sample by sample basis. Consistently within stomach samples, a population of cells with a large fraction of mitochondrial reads that had disproportionately high total counts was observed and confirmed to be made up of biologically important parietal cells; hence, appropriate mitochondrial and total count thresholds were set for these samples. For analyses where samples from different tissues were merged, whether within or across individual patients, count matrices were first normalized separately using NormalizeData. Merged normalized data was then processed with the standard Seurat pipeline of FindVariableFeatures, ScaleData and RunPCA, followed by RunUMAP and FindNeighbors with 50 principal components, and FindClusters with a resolution of 0.4 or 0.6. General tissue types were identified using established high-level markers.

Batch correction of the merged Barrett’s esophagus samples was performed with Harmony (v0.1.0) after subsetting for Barrett’s esophagus cell types following the processing approach outlined above^47^. The first round of batch correction identified more non-Barrett’s esophagus cells, as well as cells that had high rRNA expression, so the Seurat object was further subsetted. Repeating batch correction identified three more clusters of non-Barrett’s esophagus cells that were removed. Batch correction was repeated a final time, followed by UMAP and clustering on dimensions 1:50 of the Harmony embedding and a resolution of 0.8. Barrett’s esophagus cell types were identified by matching top selected genes for each cluster found using FindAllMarkers with existing annotated gastroesophageal junction and intestinal single-cell datasets^11,25^. Certain clusters were grouped together to capture a broader cell type.

The differentiation state of Barrett’s esophagus cells was determined using R package CytoTRACE (v0.3.3) with subsamplesize = 3000^48^.

### MAESTER library preparation and sequencing

Mitochondrial transcripts were enriched from the intermediate 10x cDNA libraries following the MAESTER protocol^20^. Briefly, previously amplified full-length cDNA was further amplified over six cycles in 12 separate PCR reactions containing primers that together spanned the mitochondrial transcriptome; 5-20 ng input cDNA was used in each reaction. PCR products for each sample were pooled and purified using Ampure XP (Beckman Coulter, A63881). Sequencing primer binding sites, adapters, and dual indices were added to the resulting cDNA over another six cycles of PCR and again purified using Ampure XP. Final MAESTER libraries were in the expected range of 2-100 ng ul^−1^ for concentration and fragment sizes of 300-1,500 bp. Libraries were sequenced on an Illumina NextSeq 550 with a v2.5 300 cycle kit, allocating 28 cycles for Read 1 and 270 cycles for Read 2, as well as eight cycles for both indices; a custom index 2 primer was used (10x-Ci5P).

### MAESTER sequencing data pre-processing and mapping

MAESTER raw sequencing data was demultiplexed and converted to FASTQ files using bcl2fastq (v2.20.0.422). Reads were trimmed of sequences that dropped below a quality threshold using a custom Python script: reads were broken up into 10-bp segments whose average quality (Q) score was calculated; the entire sequence following and including the first 10-bp segment with an average Q score lower than 25 was removed from the read. Reads that contained barcodes not present in the corresponding filtered 10x data for each sample were removed, and remaining Read 2 FASTQ files had the library barcode, 10x cell barcode (CB), and UMI added to the read identifier. Reads were then aligned to the 10x reference genome used above with STAR^49^.

### Mitochondrial genome variant calling

Mitochondrial mutations were called using maegatk, which was developed specifically for this protocol^20^. Before running maegatk, the CB and UMI were transferred from the Read 2 identifiers to the MAESTER BAM files as SAM tags, and the BAM files were merged with 10x mitochondrial reads. maegatk was run with the following arguments: -g rCRS-mb 100-mr 1.

### Filtering of mitochondrial variants

Allele frequencies of mitochondrial variants in single cells were calculated in R from the maegatk output and combined with accompanying variant information including mean coverage, mean Q score, and allele frequency quantiles within specified cell subsets. This information was used to select for variants within each sample that had a mean coverage of >10, mean quality of >27, and AF of >25% in at least 1% of epithelial cells and an allele frequency of >50% in at least one cell; such filtering identified germline and significant tissue-specific variants. The above filtering steps were based on previously published scripts (https://github.com/petervangalen/MAESTER-2021).

Selected variants were plotted as allele frequency heatmaps or in gene expression-derived UMAPs. In allele frequency heatmaps that looked at variants across tissues from a single patient, allele frequencies were not plotted for a variant within a tissue if that variant did not have an allele frequency of >50% in at least one cell in that tissue. Allele frequency UMAPs included cells that had a minimum coverage of 10 for the variant in question, or, in cases where it was higher, the minimum coverage was set to the mean of the mean coverage of the three main non-epithelial cell types, namely immune, fibroblast, and vascular cells. Non-epithelial cells accounted for a smaller fraction of the mitochondrial reads, so they served as a good indicator of sample sequencing depth. In order to minimize the risk of calling a cell positive for a variant as a result of ambient RNA contamination, only cells surpassing a strict allele frequency threshold of >50% that met the minimum coverage requirement were counted.

In order to confirm somatic mitochondrial variants that spanned tissue types, we developed a zero-inflated beta binomial model to capture the background noise from contamination and technical artifacts within cells for those variants. P-values could then be calculated for the variant in question in each cell to determine whether it was improbable (and therefore true) given the modeled background noise using a zero-inflated binomial test. Finally, significant cells for a given variant were identified by calculating a false discovery rate using an empirical null distribution (based on the signal in non-epithelial cells) that accounted for differences in sequencing coverage between cells. A full description of the statistical method is provided in the supplementary information.

### Tissue preparation for HCR RNA FISH

Pinch biopsies used for HCR RNA FISH were embedded fresh in OCT (Fisher, 23730571) and snap-frozen in an isopentane (O35514, Fisher)-dry ice bath, after which they were transferred to −80 °C for storage. 6-10 um tissue sections were prepared from the OCT-embedded biopsies using a cryostat (Microm) set to −20 °C and transferred to ColorFrost Plus Microscope Slides (Fisher, 12-550-17). Tissue sections were fixed in 3.7% formaldehyde (Fisher, BP531-500) solution in PBS (Invitrogen, AM9625) for 10 minutes at room temperature, followed by 2 PBS washes of 5 minutes each. 70% ethanol was used to permeabilize cells at 4 °C overnight. Slides were stored in 2xSSC (Invitrogen, AM9765) at 4 °C.

### HCR RNA FISH

Probes for target genes were designed and synthesized by Molecular Instruments. Probe sets contained between 7 and 20 probes at a concentration of 1 uM.

Slides were removed from 2xSSC and washed once with PBS before beginning the HCR RNA FISH protocol. The protocol is a modified version of the HCR v3.0 protocol that has been previously published^27,50^. In a humidified slide chamber pre-warmed to 37 °C, 200 uL of hybridization buffer (30% formamide (Invitrogen, AM9344), 10% dextran sulfate (Fisher, BP1585-100), 9 mM citric acid (pH 6.0) (Fisher, BP339-500), 50 μg mL^−1^ of heparin (Sigma-Aldrich, H5515-25KU), 1× Denhardt solution (Invitrogen, 750018), and 0.1% Tween 20 (Bio-Rad, 1610781) was added to each slide for 10 minutes. 100 uL hybridization buffer containing 0.4 pmol of each probe set was then added to the slide and covered with a glass coverslip, after which the slides were incubated for a minimum of 7 hours but up to 16 hours at 37 °C. After the probe hybridization, slides were washed three times with decreasing amounts of wash buffer (30% formamide, 9 mM citric acid (pH 6.0), 50 μg mL^−1^ of heparin, and 0.1% Tween 20) and increasing amounts of 5xSSCT for 15 minutes each, followed by 2 washes in 5xSSCT alone. Samples were then pre-amplified with 200 uL amplification buffer (10% dextran sulfate and 0.1% Tween 20) for 30 minutes at room temperature. Previously ramp-cooled (0.08 °C/s) HCR hairpins (Molecular instruments) were added to 100 uL amplification buffer at a concentration of 0.6 pmol; sections were incubated in amplification solution under a glass coverslip at room temperature concealed from light. The amplification solution was washed away with five 5-minute washes with 5xSSCT, with the last wash containing 100 ng mL^−1^ of DAPI. VECTASHIELD Vibrance antifade mounting medium (Vector Laboratories, H-1700) was applied to the sample and set under a glass coverslip for at least 4 hours at room temperature before imaging.

HCR hairpins were labeled with one of the following fluorophores: Alexa Fluor 488, Alexa Fluor 546, Alexa Fluor 594, Alexa Fluor 647, and Alexa Fluor 700.

In cases where samples were re-probed for new gene targets, coverslips and mounting medium were removed by soaking the slides in PBS at room temperature overnight. Old probes were stripped using 60% formamide in 2xSSC that was applied to the slides at 37 °C for 15 minutes. Slides were then washed three times in PBS for 15 minutes also at 37 °C. After stripping, slides were wet-mounted in 2xSSC and imaged to confirm that the probes had been successfully removed. Slides were stored in 2xSSC until the next round of HCR RNA FISH, which proceeded as described above.

### Imaging of HCR RNA FISH

HCR RNA FISH samples were imaged on an inverted Nikon Ti2-E microscope with a SOLA SE U-nIR light engine (Lumencor), an ORCA-Flash 4.0 V3 sCMOS camera (Hamamatsu), a x60 Plan-Apo λ (MRD01605) objective, and filter sets 49000 ET (Chroma), 49002 ET (Chroma), 49304 ET (Chroma), 49311 ET (Chroma), 49307 ET (Chroma), and a custom set with filters ET682.5/15x and ET725/40 (Chroma). Exposure times for the hairpin dyes were between 200 ms to 1 s, while the exposure time for DAPI was 10-20 ms.

Samples that went through subsequent rounds of HCR RNA FISH were aligned using the “Align Current ND Document” (NIS-Elements AR 5.20.02) command and converted to .tif files. The resulting files were cropped and contrasted in a custom Python script that relies on the scikit-image package to perform a gamma correction operation.

### Whole exome sequencing sample processing and library preparation

Leftover dissociated cells from the Barrett’s esophagus biopsy for patient 6 that did not go toward 10x were stored in 80% methanol, first at −20 °C for 24 hours followed by long-term storage at −80 °C; approximately 1e5 cells were saved. Total DNA was extracted from these cells following centrifugation and resuspension in 200 uL DPBS using the QIAamp DNA Mini Kit (Qiagen, 51304). 600 ng DNA were recovered, confirming our cell number estimate. Total DNA was also extracted from whole normal esophagus and gastric cardia biopsies taken from the same patient and stored in CryoStor CS10 in liquid nitrogen. Once thawed and washed in DPBS, the biopsies were chopped coarsely before being processed using the QIAamp DNA Mini Kit with the manufacturer’s recommended protocol for tissue.

Whole exome sequencing library preparation was performed using the Twist Exome 2.0 plus Comprehensive Exome Spike-in Kit (Twist Biosciences, 105036) and sequenced on an Illumina NovaSeq 6000 at the Center for Applied Genomics at the Children’s Hospital of Philadelphia with at least 100x coverage.

### Whole exome sequencing data preprocessing, somatic variant calling, and copy number analysis

Sequencing data was preprocessed following the Genome Analysis Toolkit’s (GATK) Best Practices^51^. Briefly, FASTQ files were converted to unmapped BAMs and checked for Illumina adapter sequences. Raw reads were aligned to the GRCh38 reference human genome with the Burrows-Wheeler Alignment Tool’s Maximal Exact Match algorithm (v0.7.10), after which duplicates were marked. Unless otherwise stated, the above were performed using Picard (v1.141).

Preprocessed whole exome sequencing data was analyzed with Mutect2 (GATK, v4.2.5.0) to identify short somatic mutations including single-nucleotide polymorphisms and insertions and deletions. The Barrett’s esophagus sample was run with the two matched normals, as well as the publicly available germline resource somatic-hg38_af-only-gnomad.hg38.vcf.gz (GATK). Somatic variant calls were filtered using FilterMutectCalls (GATK) and loaded into the Integrative Genomics Viewer (v2.12.3) for identification of high-quality variants in known esophageal adenocarcinoma regulators^37,38^.

Copy number alterations in the Barrett’s esophagus sample were estimated using CNVkit (v0.9.9) with the default run settings^52,53^; a pooled reference was generated from the normal esophagus and stomach samples. Copy number results were plotted directly within CNVkit using the function scatter.

### Barrett’s esophagus single-cell reference analysis of bulk transcriptomes

Count matrices of bulk transcriptomes were downloaded from the TCGA-ESCA project (https://portal.gdc.cancer.gov/projects/TCGA-ESCA). The Python package cellSignalAnalysis identified reference signals derived from the Barrett’s esophagus scRNA-seq analysis in the bulk transcriptomes, while accounting for the possibility that the mapping was incomplete^40^. Reference signals were generated from clustered scRNA-seq data by summing the raw counts for each gene across all the cells in each cluster, after which the summed counts were normalized to sum to one. cellSignalAnalysis was run using Seurat clusters that in some cases were subsequently grouped to define broader cell types. The contribution of these subclusters was combined by cell type in the output.

## Data and code availability

All code and data used in this paper will be made available upon publication.

## Acknowledgments

We thank the patients who contributed to this study. We thank all members of the Shaffer Lab for their valuable feedback and support. We thank L. Bugaj, P. Cámara, A. Raj, and K. Zaret for insightful comments on the manuscript. We thank C. Ren and D. Dykes for assistance with illustrations. We thank L. Dolinsky, G. Park, and A. Benitez for assistance with sample retrieval. We thank R. Wu and A. Sahasrabuddhe, as well as the Genomics Facility at the Wistar Institute, for assistance with sequencing. We thank the Center for Applied Genomics at the Children’s Hospital of Philadelphia for assistance with whole exome library preparation and sequencing. R.A.R.H. acknowledges support from the NSF Graduate Research Fellowship (DGE-1845298). A.B.M. acknowledges support from NIH grant R01DK124266. G.W.F acknowledges support from the Center for Molecular Studies in Digestive and Liver Diseases (NIH/NIDDK P30 DK050306). S.M.S. acknowledges support from the NIH Director’s Early Independence Award DP5OD028144, a donation to the Institute for Regenerative Medicine at the University of Pennsylvania from Larry and Mickey Magid, and the Institute for Translational Medicine and Therapeutics of the Perelman School of Medicine at the University of Pennsylvania (NIH NCATS UL1TR001878).

## Author contributions

Conceptualization, R.A.G. and S.M.S.; Methodology, R.A.G. and S.M.S.; Investigation, R.A.G. and R.A.R.H.; Software, R.A.G., R.A.R.H., and J.R.; Formal Analysis, R.A.G., J.R., and N.R.Z.; Data Curation, R.A.G. and R.A.R.H.; Writing – Original Draft, R.A.G and S.M.S.. Writing – Review and Editing, R.A.G. and S.M.S.; Visualization, R.A.G., R.A.R.H., and S.M.S.; Resources, M.D., T.A.K., A.B.M, G.W.F; Funding Acquisition, S.M.S.; Supervision, N.R.Z. and S.M.S.

