## Supplementary Figures for "Clonal cell states link Barrett’s esophagus and esophageal adenocarcinoma"

**R. A. Gier et al.**

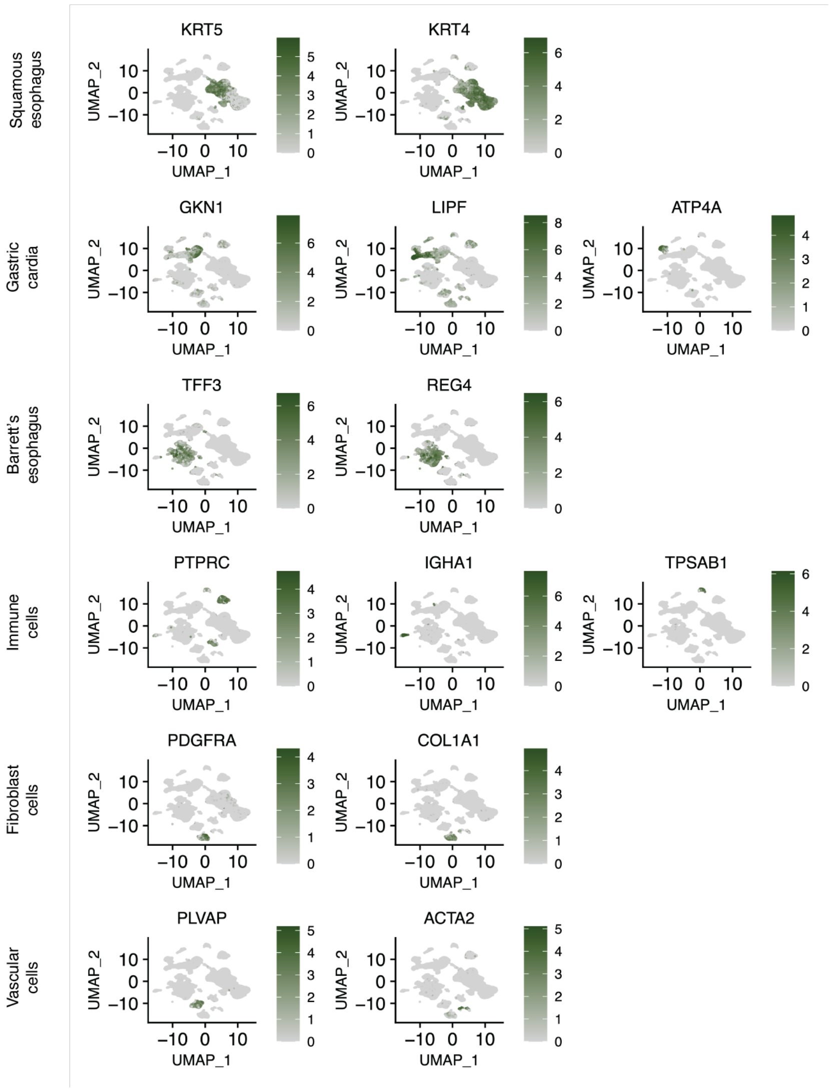


**Fig. S1. Annotation of GEJ tissues and supporting cell populations.** UMAPs of scRNA-seq of all the samples collected in this study featuring expression of high-level tissue markers of the normal esophagus (*KRT5*, *KRT4*), the gastric cardia (*GKN1*, *LIPF*, *ATP4A*), Barrett's esophagus (*TFF3, REG4*), immune cells (*PTPRC*, *IGHA1*, *TPSAB1*), fibroblasts (*PDGFRA*, *COL1A1*), and vascular cells (*PLVAP*, *ACTA2*).

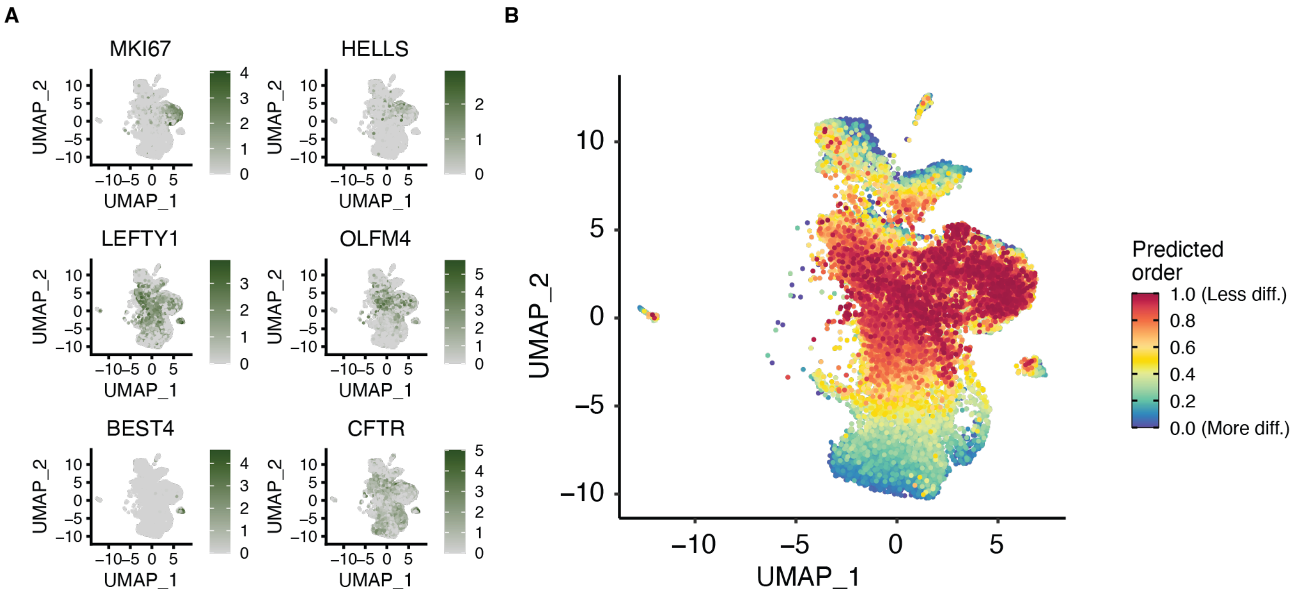


**Fig. S2. Markers of proliferative and immature Barrett's esophagus cell types and their differentiation state.** (**A**) UMAPs of scRNA-seq of consensus Barrett's esophagus cells featuring the expression of key proliferative (*MKI67*, *HELLS*), stem (*LEFTY1*, *OLFM4*), and rare cell (*BEST4*, *CFTR*) markers. (**B**) UMAP of scRNA-seq of consensus Barrett's esophagus cells colored by CytoTRACE analysis results, where cells are ordered along a differentiation continuum.

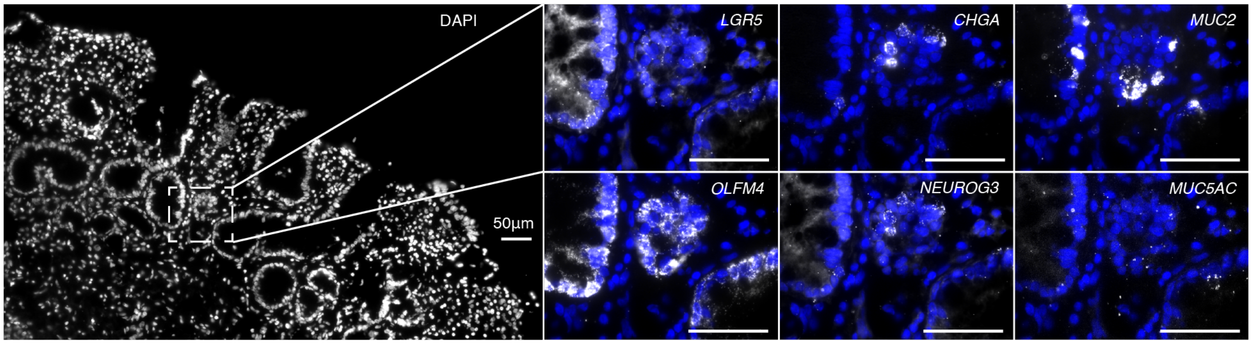


**Fig. S3. Spatial location of Barrett's esophagus secretory progenitors.** Multiplexed HCR RNA FISH of fresh frozen Barrett's esophagus sections for *LGR5* (stem), *OLFM4* (stem), *CHGA* (enteroendocrine), *NEUROG3* (enteroendocrine progenitor), *MUC2* (goblet), and *MUC5AC* (foveolar) with DAPI counterstain. Secretory progenitor cells co-express stem cell marker *OLFM4* and either *CHGA* and *NEUROG3* or *MUC2*, suggesting that OLFM4 progenitors are direct precursors of these more mature cell types. Scale bars, 50 µm.

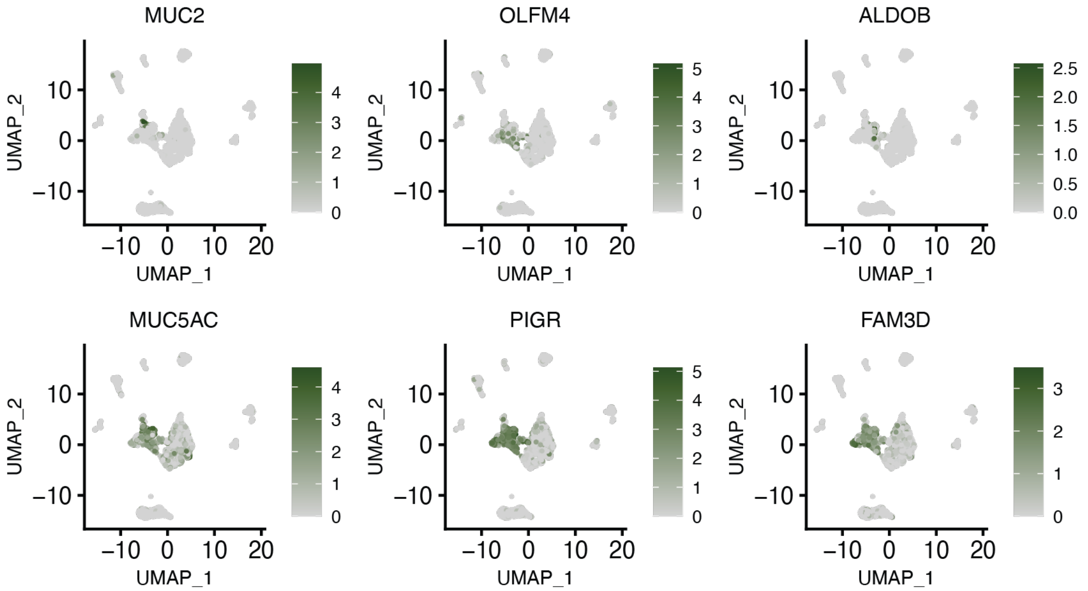


**Fig. S4. Markers of typical Barrett's esophagus cell types and barrier integrity genes in dysplastic biopsy.** UMAPs of scRNA-seq of cells from a biopsy taken from a patient 6 with high-grade dysplasia examined in depth in Fig. 4 featuring typical Barrett's esophagus cell type markers and barrier integrity genes: *MUC2* (goblet), *OLFM4* (stem), *ALDOB* (enterocyte), *MUC5AC* (foveolar), *PIGR* (barrier), and *FAM3D* (barrier).

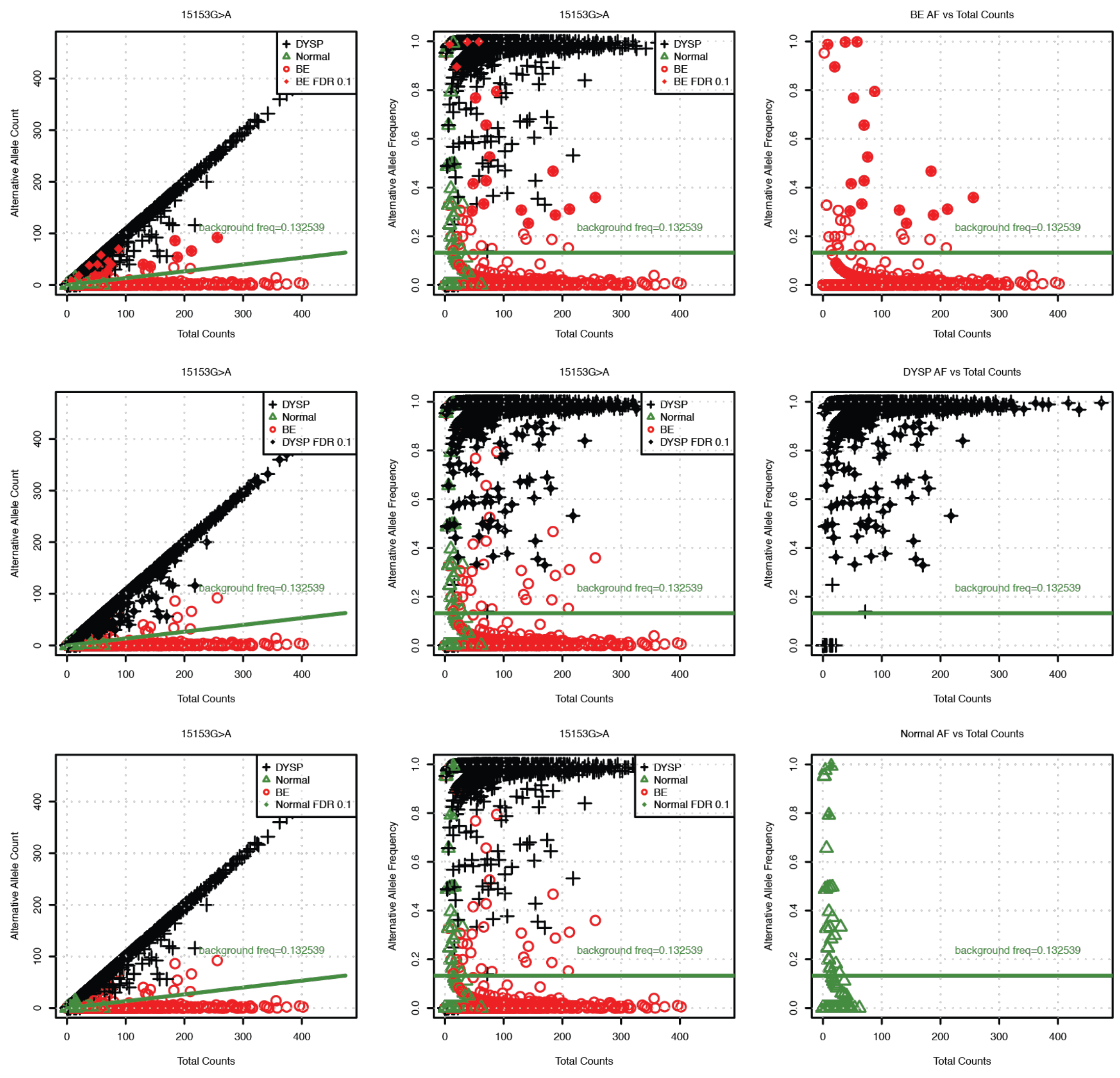


**Fig. S5. ZIBB model applied to 15153G>A in Barrett's esophagus, dysplastic, and non-epithelial cell populations.** Allele frequency profiles of mitochondrial variant 15153G>A in the cells from the Barrett's esophagus biopsy from patient 6, plotted with the estimated background contamination rate in green. The first column of plots shows the read counts for the alternative allele versus the total read counts at that position in all cells. The second column of plots shows the allele frequency versus the total read counts in all cells. The third column of plots is the same as the second column, only featuring a single cell type. Each row of plots highlights one of three cell types, Barrett's esophagus (BE) in the first, dysplasia (DYSP) in the second, and non-epithelial cells (Normal) in the third; highlighted cells were determined to be significantly different from background by the ZIBB model. Of note are the 18 Barrett's esophagus cells with the 15153G>A variant, of which one cell is expected to be a false positive based on the separate-class FDR.

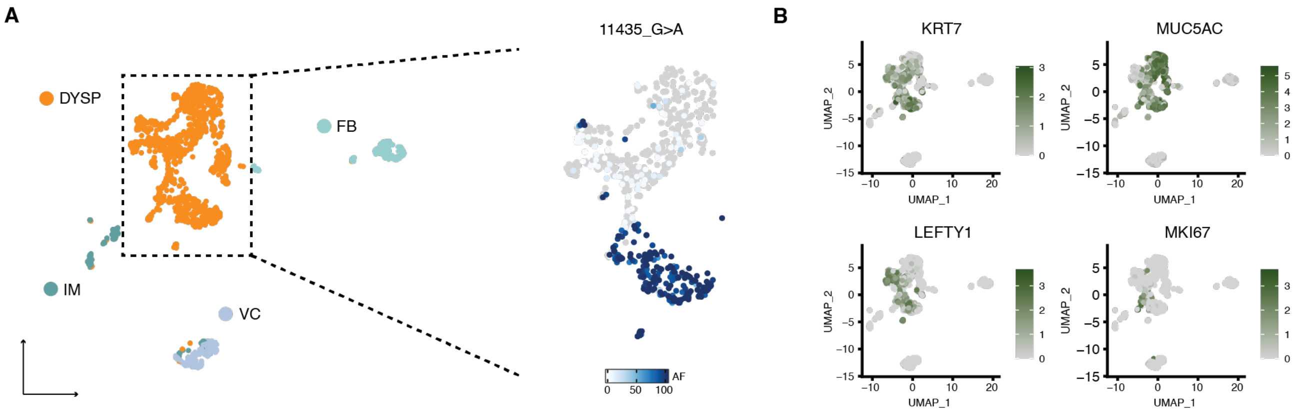


**Fig. S6. Clonal and molecular characterization of second dysplastic case.** (**A**) UMAP of scRNA-seq of a single biopsy from patient 7, who was diagnosed with low-grade dysplasia, with the major tissue types labeled. The callout of the dysplastic cells is colored by the allele frequency of the mtDNA mutation 11435G>A. (**B**) UMAP from (A) featuring the expression of differentially expressed genes within the dysplastic cells.

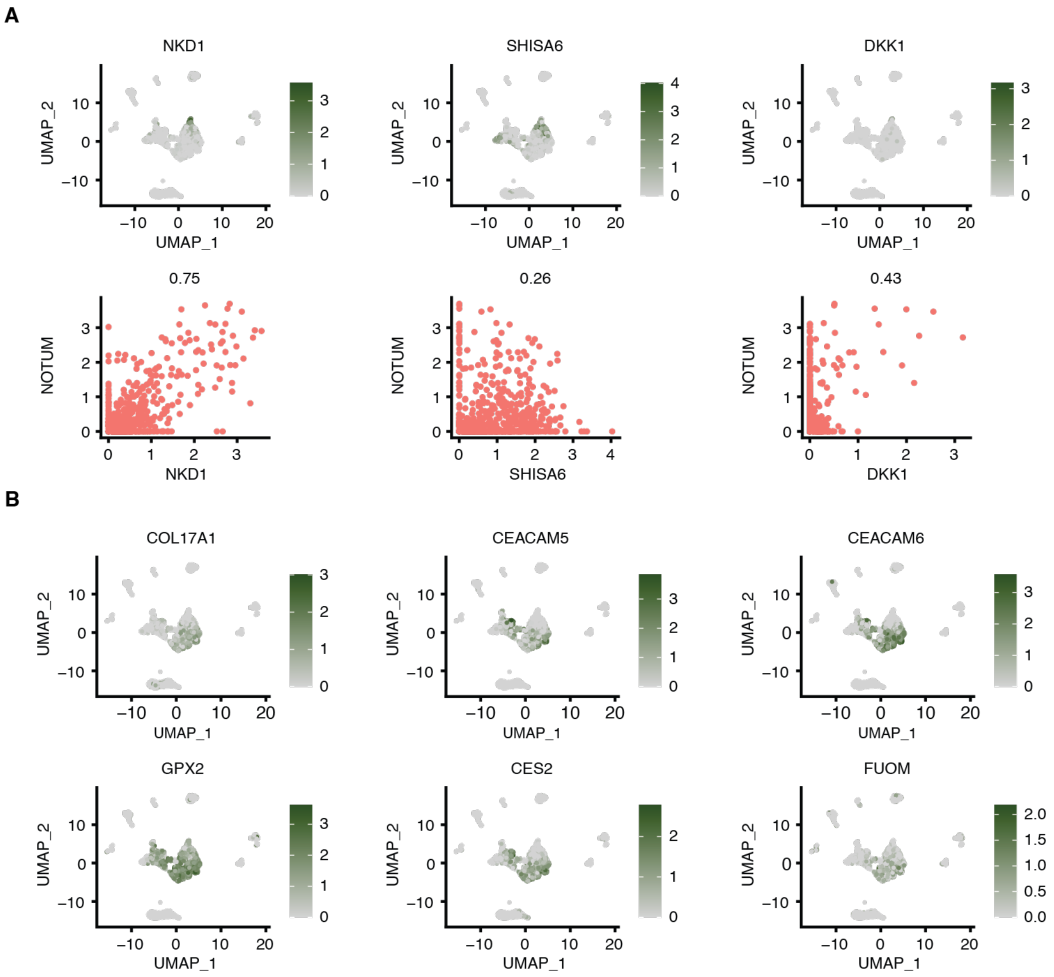


**Fig. S7. Differential expression of WNT antagonists and potential fitness genes within dysplastic clones.** (**A**) UMAPs of scRNA-seq of cells from a biopsy taken from a patient 6 with high-grade dysplasia examined in depth in Fig. 4 featuring differential expression of *WNT* antagonists within the dysplastic cells. Below the UMAPs are scatter plots of each gene with NOTUM, where the Pearson correlation coefficient is shown above. (**B**) UMAPs of the same sample as in (A) featuring the expression of genes differentially expressed in the dysplastic subclone marked by mtDNA mutation 5215T>C (see Fig. 4C).
